# Beneficial effects of two Ayurvedic formulations, *Saraswata Ghrita* and *Kalyanaka Ghrita* on survival and on toxic aggregates in *Drosophila* models of Huntington’s and Alzheimer’s disease

**DOI:** 10.1101/2021.10.06.463232

**Authors:** Swati Sharma, Girish Singh, Kishor Patwardhan, Subhash C. Lakhotia

**Affiliations:** Department of Kriya Sharir, Faculty of Ayurveda, Institute of Medical Sciences, Banaras Hindu University, Varanasi 221005, India; Centre of Biostatistics, Institute of Medical Sciences, Banaras Hindu University, Varanasi 221005, India; Cytogenetics Laboratory, Department of Zoology, Institute of Science, Banaras Hindu University, Varanasi 221005, India

**Keywords:** Dementia, Neurodegeneration, Amyloid aggregates, PolyQ Aggregates

## Abstract

In order to understand the health promotive, rejuvenative and disease preventive approach of the Ayurvedic system of medicine in the light of current principles, we examined two Rasayana formulations, viz., *Kalayanaka Ghrita* (*KG*) and *Saraswata Ghrita* (*SG*) for their effects in Alzheimer’s (AD) and Huntington’s (HD) neurodegenerative disease models of *Drosophila*. Initial experiments involving feeding of wild type flies on food supplemented with 0.05%, 0.25% and 0.5% (w/v) *KG* or *SG* revealed 0.05% to be without any adverse effect while higher concentrations caused dose-dependent reduction in pupation frequency and adult life span in wild type flies. Rearing *GMR-GAL4>127Q* (HD model) and *ey-GAL4>Aβ42* (AD model) larvae and adults on 0.05% or 0.25% *SG* or *KG* supplemented food enhanced the otherwise significantly reduced larval lethality and enhanced their median life span, with the 0.25% *SG* or *KG* concentrations being less effective than the 0.05%. In parallel with the better larval survival and enhanced adult life span, feeding the HD and AD model larvae on either of the *Ghrita* supplemented food (0.05% and 0.25%) substantially reduced the polyQ aggregates or amyloid plaques, respectively, in the larval eye discs. The present first in vivo organismic model study results have clinical implications for the increasing burden of age-associated dementia and neurodegenerative diseases like AD and HD in human populations.

## 1. Introduction

Dementia is a group of age-associated and other specific medical conditions due to some abnormal brain changes that impair the person’s cognitive ability and hamper daily living (Jameson, 2018). Some of the major clinical conditions are Alzheimer’s disease (AD), vascular dementia, Lewy body dementia and frontotemporal dementia. In addition, Parkinson’s disease, Huntington’s disease (HD), Progressive supra nuclear palsy, Creutzfeldt-Jakob disease, and other conditions like prion disease, neurosyphilis and any traumatic encephalopathy also cause dementia (Jessen et al., 2020).

AD, contemporarily the most common cause of neurodegenerative disorders, is associated with progressive decline of memory and cognition (Albert et al., 2011). Characteristic pathology of AD includes, besides the progressive memory loss, accumulation of Amyloid-β (Aβ) plaques, hyper-phosphorylated Tau protein based neurofibrillary tangles, and synaptic loss particularly due to deficiency of acetylcholine and consequent degeneration of cholinergic neurons in cortex, hippocampus etc. The earliest pathology is perhaps the oxidative stress, leading to accumulation of Aβ plaques and neuro-fibrillary tangles (Nunomura et al., 2001). Currently it has become the most significant challenge that affects social, medical as well as economic sector (Polis and Samson, 2019).

HD is a progressive neurodegenerative disorder, associated with chorea, progressive cognitive impairment and behavioral issues, and belongs to a class of inherited neurological diseases caused by expansion of the CAG-nucleotide repeats, leading to extensive polyQ (poly-glutamine) tracts in their respective protein products (Zoghbi and Orr, 2000; Mallik and Lakhotia 2010). Proteins with expanded polyQ tracts form cytoplasmic and nuclear protein aggregates which are preferentially toxic to neurons, e.g., the medium spiny neurons of striatum and certain subsets of neurons in the cortex, leading to neurodegeneration and chorea (Beal et al., 1993).

Dementia is a common feature in both AD and HD. AD affects storage and/or retrieval of semantic memory (Rohrer et al., 1999). Alzheimer’s condition is also suggested to contribute to the cognitive decline in elderly Huntington’s patients (Davis et al., 2014).

In view of the severe physical, mental, emotional and financial stress faced by dementia patients, their families and caregivers, many therapeutic applications have been developed (Jessen et al., 2020; Livingston et al., 2020). However, most of these offer only marginal benefits since they mostly target symptoms rather than the root cause/s. Further, reversal of cognitive decline through such therapies in some cases is also reported to be associated with other complications (Pickett et al., 2018). In this context, use of complementary, traditional and alternative medicine approaches may be more effective (Lakhotia, 2019; Sharma et al., 2021).

The potential role of indigenous traditional knowledge systems in preventing and treating various clinical conditions is gaining wide acceptance and approval by the World Health Organization (WHO, 2013). *Ayurveda* is one of the oldest and widely practiced traditional systems of healthcare in India, Sri Lanka, Nepal and other regions in South Asia. In order to provide effective health care to populations, the societies must consider integrating the available traditional knowledge systems with the different prevalent practices (Lakhotia 2020).

*Ayurveda* prescribes individualized lifestyle, dietary and other interventions for prevention and management of various health conditions. Among the available traditional approaches, the *Rasayana* group of preparations hold a high potential for treating cognitive impairments. The mostly poly-herbal *Rasayana* formulations are prepared using well-defined complex preparatory processes. Many of the component herbs or their ‘active principle molecules’ have been evaluated for their antioxidant, free radical-scavenging and neuro-protective properties through experimental, preclinical and clinical studies (Farooqui et al., 2018). *Ashwagandha, Brahmi, Shankhpushpi, Madukparni, Jatamansi* have been studied for their nootropic properties (Nahata et al., 2008; Russo et al., 2001;Begum et al., 2008; Stough et al., 2008; Matsuda et al., 2001; Ganguli et al., 2000; Uabundit et al., 2010; Sultana et al., 2005; Kumar et al., 2012). However, the Ayurvedic approach typically involves administration of *Rasayana* that contain multiple herbs and/metals or other components processed through various pharmaceutical steps to produce the given formulation. Therefore, rather than evaluating individual components, assessing efficiency of *Ayurvedic* preparations in the form in which they are prescribed in traditional practice is a better approach to ascertain their potential in treatment and the mechanisms underlying their therapeutic applications (Lakhotia 2019, 2020).

In the present study, we used *Saraswata ghrita* (*SG*) and *Kalyanaka ghrita* (*KG*), which are commonly prescribed ghee-based interventions for dementia, AD and HD conditions. *Ashtanga Hridaya*, a classical Ayurveda textbook composed by Vagbhata around 6^th^ Century AD, also indicated these formulations for persons with loss of memory, dementia and impairment of higher intellectual functions. (Murthy, 2009). In the present study, we examined biological effects of *SG* and *KG* in *Drosophila melanogaster* (fruit fly) model. In recent years, the *Drosophila* fly model has emerged as a very powerful model for health-related issues since many human diseases, especially the diverse neurodegenerative disorders, have been modelled in flies to understand the molecular bases of their pathology and identification of the affected gene/s, which opens the possibilities of developing new therapeutics (Bilen and Bonnini, 2005; Mallik and Lakhotia, 2010; Pussing et al, 2013; McGurk et al., 2015); Perrimon et al 2016; Chow and Reiter, 2017; Oriel and Lasko 2018; Bellen et al 2019). Earlier studies (Dwivedi et al, 2012; 2013, 2015) have established the fly as a good model for examining effects of various Ayurvedic *Rasayanas* at organismic, tissue and molecular levels. Results of the present study show that at low dosages, neither of these *ghrita* formulations have any adverse effect on life-history parameters of the organism. More significantly, both these *ghritas* substantially suppress the polyQ inclusion bodies and the amyloid plaques in HD and AD transgene expressing tissues, respectively. This is associated with significant improvement in survival of the afflicted individuals. It is also notable that while higher dietary concentrations of *KG* and *SG* are deleterious for wild type flies, they provide protective advantages to organisms expressing AD or HD symptoms.

## 2. Materials and Methods

### 2.1 Rasayana Formulations

*Saraswata ghrita* and *Kalyanaka ghrita*, the two formulations used in this study, were obtained from the Arya Vaidyasala, Kottakkal (India). Their contents are shown in Table 1.

**Table I.** Components of *Kalyanaka Ghrita* and *Saraswata Ghrita* as per Ayurvedic Pharmacopoeia of India, Department of AYUSH, GOI for 10g of final preparation

| *Kalyanaka Ghrita* |  | *Saraswata Ghrita* |  |
| --- | --- | --- | --- |
| Component (Botanical name) | Quantity | Component (Botanical name) | Quantity |
| **Ghrita** or *Ghee* | 12.723g | **Ghrita** or *Ghee* | 11.826 g |
| **Amalaki** (*Phyllanthus emblica*) | 0.092g | **Ksheer** or Cow Milk | 11.130 ml |
| **Bhadraila** (*Amomum subulatum*) | 0.092g | **Haritaki** (*Terminalia chebula*) | 0.174g |
| **Brihati** (*Solanum anguivi*) | 0.092g | **Maricha** (*Piper nigrum*) | 0.174g |
| **Dadima** (*Punica granatum*) | 0.092g | **Patha** (*Cyclea peltata*) | 0.174g |
| **Danti** (*Baleospermum montanum*) | 0.092g | **Pippali** (*Piper longum*) | 0.174g |
| **Daruharidra** (*Berberis aristate*) | 0.092g | **Saindhava** or Rock salt | 0.174g |
| **Devadaru** (*Cedrus deodara*) | 0.092g | **Shunthi** (*Zingiber officinale*) | 0.174g |
| **Ela** (*Elettaria cardamomum*) | 0.092g | **Shigru** (*Moringa oleifera*) | 0.174g |
| **Ervaruka** (*Prunus avium*) | 0.092g | **Vacha** (*Acorus calamus*) | 0.174g |
| **Haridra** (*Curcuma longa*) | 0.092g |  |  |
| **Haritaki** (*Terminalia chebula*) | 0.092g |  |  |
| **Hima** (*Santalum album*) | 0.092g |  |  |
| **Kushtha** (*Saussurea costus*) | 0.092g |  |  |
| **Maltimukula** (*Jasminum grandiflorum*) | 0.092g |  |  |
| **Manjishtha** (*Rubia cordifolia*) | 0.092g |  |  |
| **Nagakeshara** (*Mesua ferrea*) | 0.092g |  |  |
| **Nata** (*Valeriana jatamansi*) | 0.092g |  |  |
| **Padmaka** (*Prunus cerasoides*) | 0.092g |  |  |
| **Phalini** (*Callicarpa macrophylla*) | 0.092g |  |  |
| **Prishnaparni** (*Desmodium gangeticum*) | 0.092g |  |  |
| **Sariva** (*Hemidesmus indicus*) | 0.092g |  |  |
| **Shalaparni** (*Pseudarthria viscida*) | 0.092g |  |  |
| **Talishpatra** (*Abies spectabilis*) | 0.092g |  |  |
| **Utpala** (*Kaempferia rotunda*) | 0.092g |  |  |
| **Utpalasariva** (*Ichnocarpus frutescens*) | 0.092g |  |  |
| **Vibhitaki** (*Terminalia bellirica*) | 0.092g |  |  |
| **Vidnaga** (*Embelia ribes*) | 0.092g |  |  |
| **Vishala** (*Citrullus colocynthis*) | 0.092g |  |  |

#### 2.1.1 Preparatory procedure

The standard method of preparation of both these formulations, as followed at the Arya Vaidyasala, involved washing of the specified quantities of herbal ingredients, following which they were dried and reduced to coarse powder with an electrically operated Chopper/ pulverizer. The pulverised herbal combination was ground in electrically operated Grinding Stone with the required quantity of water added to obtain a pasty material, the *kalka*. The *kalka* was then transferred, together with the specified quantity of ghee, to the processing pan, a double walled vessel with SS-304 contact surface, for cooking by steam which passes through the space between the double walls, so that the skin temperature is not more than 100^°^C. During the steam heating, required quantity of water was gradually added to the mixture in the pan as the boiling started, which was continued till the *kalka* attained the specified *madhyamapaka* character. Immediately thereafter, the material was filtered through fine cloth filter to collect the processed ghee. The solid debris remaining on the filter was compressed using an electrically operated hydraulic press to squeeze out the remaining ghee. This was added to the processed ghee and allowed to cool naturally.

#### 2.1.2 *Drosophila* rearing and stocks

*Drosophila melanogaster* stocks were maintained under uncrowded condition at 24°C±1°C. Flies were fed on standard fly food containing agar, maize powder, yeast and sugar. The following fly stocks were used.

i. Oregon R^+^ **-** Wild type strain of *Drosophila melanogaster*.
ii. *w*^*1118*^; *UAS-Aβ*^*H32*.*12*^ *ey-GAL4/CyO* stock was used for Alzheimer’s model. The *UAS-Aβ*^*H32*.*12*^ tran*sg*ene is referred to here as *UAS-Aβ42*. The *UAS-Aβ42 ey-GAL4* larvae show accumulation of amyloid protein aggregates in their eye disc cells since the *ey-GAL4* drives expression of the *UAS-Aβ42* transgene in developing larval eye discs posterior to the morphogenetic furrow to produce the mutant Aβ42 protein. The collected 1^st^ instar larvae were mostly *UAS-Aβ42 ey-GAL/CyO* since all the *CyO* homozygotes die as embryo, while very few *UAS-Aβ42 ey-GAL4* homozygotes survive to late larval stage. This genotype has been widely used as an AD model since it mimics various symptoms of AD, including the characteristic age dependent cognitive decline (Finelli et al 2004; Iijima et al 2004).
iii. *w*^*1118*^; *GMR-GAL4 UAS-127Q/CyO-GFP* **–** The *GMR-GAL4* driven expression of the *UAS-127Q* transgene leads to accumulation of HA-tagged polyQ inclusion bodies (IBs) or polyQ aggregates posterior to the morphogenetic furrow in late 3rd instar larval eye discs (Davies et al., 1997; Klement et al., 1998). In this case also, most of the collected 1^st^ instar larvae were *GMR-GAL4 UAS-127Q/CyO-GFP* since while all the *CyO-GFP* homozygotes die as embryo, very few *GMR-GAL4 UAS-127Q* homozygotes survive to late larval stage. The accumulation of the IBs disrupts the ommatidial units and leads to the irregular axonal projections from the rhabdomeres to the optical lobe of the brain (Mallik and Lakhotia, 2010). Consequently, in addition to the extensive presence of IBs, disarrayed rhabdomeric complexes and irregularly arrayed or missing axons are characteristic features in eye discs of *GMR-GAL4>UAS-127Q* larvae reared on normal food.

#### 2.1.3 Administration of the ghrita Rasayanas to Drosophila

The desired quantity of the *ghrita Rasayana* was mixed with the freshly prepared fly food just before its solidification following the method described earlier (Dwivedi et al 2012). The concentrations (weight of *ghrita* formulation/volume of cooked fly food prior to solidification) of *KG* or *SG* in the food were 0.05%, 0.25%, or 0.5%. After thorough mixing, the food was poured into fly culture bottles, vials or petri plates, as required and allowed to cool and solidify before use. In the treatment sets, the larvae and flies were allowed to feed continuously on the formulation-supplemented food. Parallel controls were kept with larvae and flies reared on normal food without the *Rasayana* supplement. For each experiment, the control and the formulation supplemented food bottles, vials/petri dishes were prepared from the same batch of food; likewise, all larvae/adults for a given experiment were derived from a common pool of eggs of the desired genotype and reared in parallel on the control or formulation supplemented food. Eggs for all experiments were collected from flies that had not been exposed to either of these formulations in their lives.

### 2.2 Assessment of effects of the formulation feeding on life history traits in wild type flies

#### 2.2.1. Larval and pupal development assay

Synchronized freshly hatched 1^st^ instar larvae of the desired genotype were collected and transferred to regular (control) or the *ghrita Rasayana* supplemented food vials. Total number of larvae that pupated were recorded in each case. The number of pupae that hatched as flies emerging from these pupae were also recorded and expressed as % of the original numbers of eggs that were followed through larval and pupal development in each case.

#### 2.2.2. Lifespan assay

This was carried out by rearing freshly eclosed flies, as above, on regular or one of the *ghrita Rasayana* supplemented food since the 1^st^ instar larval stage. After every 2-3 days, dead flies in the given bottle were recorded and the surviving flies were transferred, without anaesthetization, to fresh bottles with respective control or supplemented food. This was repeated till 60 days after eclosion. The median life span, estimated by identifying the day when 50% of the original sample size is dead, was calculated for each set.

All experiments were carried out at 24±1° in four replicates, with 100 wild type or 80 disease model 1^st^ instar larvae as the starting material in each.

### 2.3 Immunostaining

Eye imaginal discs from late third instar larvae of the desired genotypes (see Results) were dissected out, fixed and immunostained as described earlier (Prasanth et al, 2000). The following primary and secondary antibodies were used as required. The *Anti-HA* rabbit polyclonal antibody (Y-11, sc-805, Santa Cruz, USA), raised against a peptide of the influenza haemagglutinin (HA) protein, was used for immunostaining of the HA-tagged 127Q polypeptide at a working dilution of 1:40. The *mAb22C10* mouse monoclonal antibody (Developmental Studies Hybridoma Bank, Iowa) was used to stain the peripheral neurons and a subset of the central nervous system neurons at 1:100 working dilution. The Anti-Amyloid Peptide β (A1976-25UG, Sigma-Aldrich, India), raised in rabbit, was used to immunostain the mutant Aβ amyloid plaques at a working dilution of 1:100.

The different secondary antibodies used were: *Alexa Flour 488 donkey anti-rabbit IgG* (Molecular Probes, USA) at a dilution of 1:200 to detect the anti-HA primary antibody raised in rabbit; *Cy3 conjugated anti-mouse IgG* (Whole molecule; Sigma-Aldrich, India) at 1:200 working dilution to detect the *mAb22C10* primary antibody raised in mouse and *Alexa Flour 546 donkey anti-rabbit IgG* (Molecular probes, USA) at a dilution of 1:200 to detect the anti-Aβ42 primary antibody.

Nuclei were stained with DAPI (4’, 6-diamidino-2-phenylindole dihydrochloride, Sigma-Aldrich, India) at 1μg/ml.

A minimum of 8-10 eye discs for each genotype and treatment were processed for immunostaining and microscopy and each set was replicated at least twice.

### 2.4 Microscopy and image analysis

Confocal imaging was carried out with LSM510 Meta Zeiss laser scanning confocal microscopy using appropriate laser, dichroic and barrier filters. All the images were assembled using Adobe Photoshop CS 8.0.

### 2.5 Statistical analysis

SigmaPlot 11.0 and SPSS 16.0 software were used for statistical analyses. All percentage data were subjected to arcsine square root transformation. One-way ANOVA and Dunnett’s post-hoc test were applied to compare the means. Data are expressed as mean ± S.E. of mean (SEM) of the replicates.

## 3. Results

### 3.1 Effects of feeding wild type larvae and adults on different concentrations of *Saraswata Ghrita* (*SG*) or *Kalyanaka Ghrita* (*KG*) on life history parameters

The percentage of the 1^st^ instar larvae that developed to pupal stage and finally emerged as adult flies and the median life span of the emerging adult flies were taken as indicative parameters. As the data in Fig. 1A show, the lowest concentration (0.05% w/v) of *KG* or *SG* did not affect the pupation and fly emergence or the median life span of wild type flies, while feeding on higher concentrations (0.25% and 0.5% w/v) of *SG* or *KG* during larval and adult stages significantly reduced the median life span of adult flies; the frequency of pupation was also affected when the wild type larvae were reared on 0.5% SG, but most of those that pupated emerged as flies (Fig. 1). Concentrations higher than 0. 5% (0.75%and 1.0%) of the *SG* or *KG* showed dose-dependent greater reduction in larval survival and adult life span (data not presented). Therefore, only 0.05% and 0.25% *KG* or *SG* supplemented food were examined for their effects in the HD and AD *Drosophila* models.

**Fig. 1.**
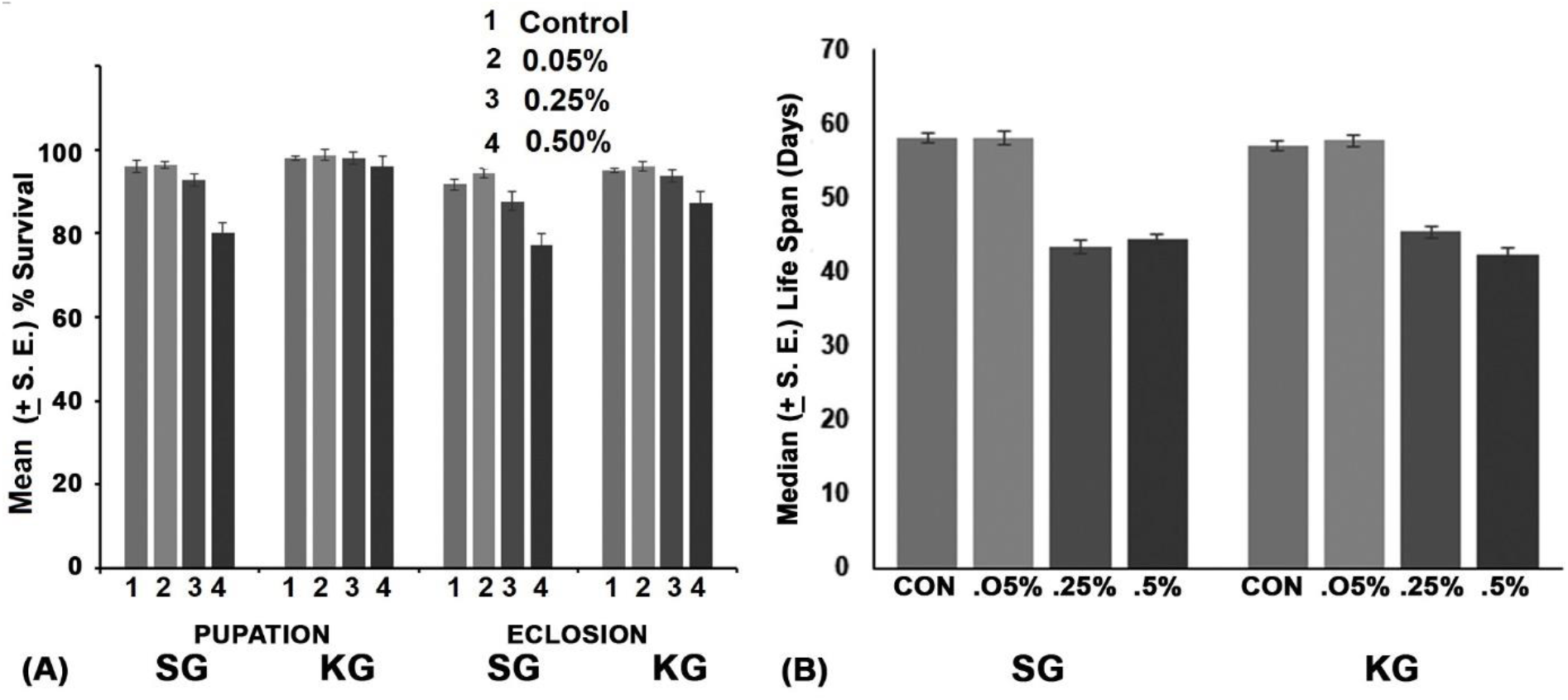
**Effects of feeding wild type (Oregon R) larvae and flies on *Saraswata Ghrita* (*SG*) or *Kalyanaka Ghrita* (*KG*) supplemented food on (A) mean % (± S. E**., **Y-axis) survival till pupation and eclosion (X-axis), and (B) median (± S. E**., **Y-axis) life span of adults. Concentrations of the respective *Ghrita* used are indicated on upper right corner in (A) and on X-axis in (B)**.

### 3.2 Effects of feeding larvae and adults on different concentrations of *Saraswata ghrita* (*SG*) or *Kalyanaka ghrita* (*KG*) on life history parameters in HD and AD flies

Data presented in Fig. 2 show that, in agreement with earlier results (Dwivedi et al, 2013), the *GMR-GAL4>UAS-127Q* individuals reared on normal food show significant death during the larval stage while the adults have a greatly reduced median life span (∼26 days) compared to ∼57-58 days of Oregon R flies (Fig. 1 and 2). Interestingly, feeding these larvae on either of the *ghrita* supplemented food at 0.05% concentration significantly improved their emergence as adult (Fig. II A) as well as the median life span of the emerged flies (Fig. 2B). In each case, the improvement in fly emergence and life span at 0.25% concentration was less than that with 0.05%, although significantly more than the respective values for those reared on un-supplemented control food (Fig. 2A).

**Fig. 2.**
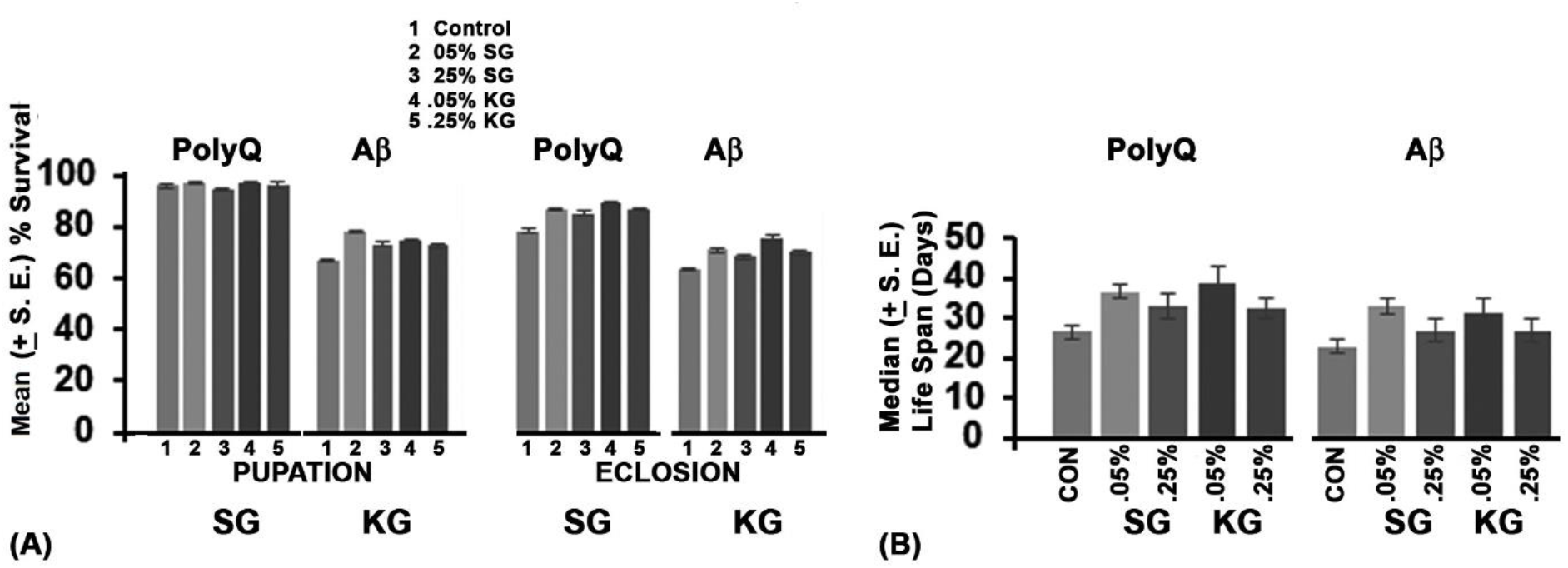
**Effects of feeding polyQ (*GMR-GAL4>UAS-127Q*) or Aβ42 (*ey-GAL4>UAS-Aβ42*) expressing larvae and flies on *Saraswata ghrita* (*SG*) or *Kalyanaka ghrita* (*KG*) supplemented food on (A) mean % (± S. E**., **Y-axis) survival till pupation and eclosion (X-axis), and (B) median (± S. E**., **Y-axis) life span of adults. Legend to the concentrations of the *ghrita* used are indicated on upper part in (A) while these are indicated on X-axis in (B)**.

Compared to the Oregon R wild type, the *ey-GAL4>UAS-Aβ42* AD larvae reared on normal food showed significantly high larval and pupal lethality with reduced median life span of adults. All these parameters were affected more than in the case fo polyQ expressing larvae/flies reared on normal food (Fig. 2). Similar to the effects of *KG* and *SG* dietary supplement in the HD model, the incidence of successful pupation and emergence, and the median life span of *ey-GAL4>UAS-Aβ42* individual were also enhanced when reared on *SG* or *KG* supplemented food, with their lower (0.05%) concentration being more effective than the higher concentration (Fig. 2).

### 3.3 The improved survival of HD and AD model larvae reared on *KG* or *SG* supplemented food is associated with substantial suppression of the toxic aggregates in *127Q* or *Aβ42* expressing eye disc cells

The above results on life-history parameters indicated that both formulations seem to partially mitigate the systemic damage inflicted by expression of either the expanded polyQ protein or the mutant Aβ protein in eye discs. Their expression results in accumulation of aggregates of the mutant proteins, which are believed to be primarily responsible for the toxicity and neurodegeneration. Therefore, eye discs of late third instar larvae in which expression of the mutant proteins were targeted, immunostained and examined by confocal microscopy to assay the levels of the polyQ inclusion bodies (IBs) or the mutant Aβ protein amyloid aggregates in the respective model eye discs.

Since the 127Q protein carries the HA-tag, the eye discs from *GMR-GAL4>UAS-127Q* larvae were immunostained with anti-HA antibody to detect the polyQ aggregates. The axons of the ommatidial units were visualized by immunostaining with the mAb22C10 which specifically decorates axons and their cell bodies. As seen in Fig. 3A-A”, the eye disc cells posterior to the morphogenetic furrow in the 127Q expressing larvae reared on normal food have massive accumulation of polyQ inclusion bodies and grossly disrupted organization of optical neurons and their axons. Interestingly, the polyQ IBs were substantially reduced in eye discs from *GMR-GAL4>UAS-127Q* larvae reared on *KG* or *SG* supplemented food (fig. 3B-E”). In parallel with the reduced load of polyQ aggregates in the formulation fed larval eye discs, the integrity of axonal projections, as revealed by the mAb22C10 antibody staining, was also significantly improved.

**Fig. 3.**
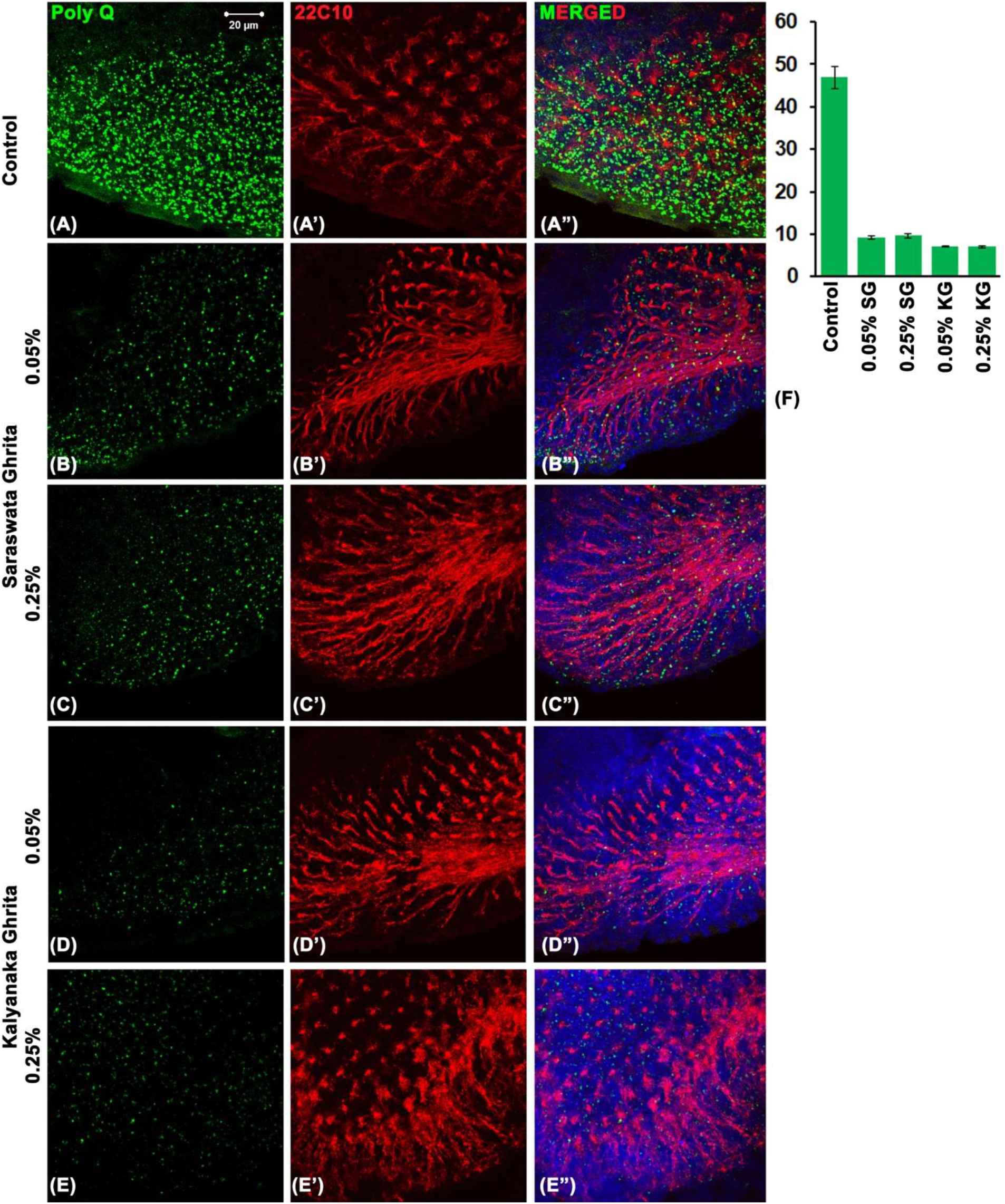
Rearing on *Saraswata ghrita* or *Kalyanaka ghrita* supplemented food substantially reduced accumulation of polyQ aggregates (green) in *GMR-GAL4>127Q* eye discs. **(A-E”)** are confocal projection images of *GMR-GAL4>127Q* eye discs stained for polyQ protein (green, A-E) and neuronal cells and axons (red, 22C10, **A’-E’**); the column **A”-E”** shows corresponding merged images including DAPI-stained nuclei (blue). The specific feeding regimes are noted on left side of the rows. (**F**) Histograms of mean (± S.E.) fluorescence intensity of polyQ immunostaining (expressed in arbitrary units, Y-axis) in imaginal discs from larvae reared on control or *Saraswata Ghrita* (*SG*) or *Kalyanaka Ghrita* (*KG*) supplemented food (concentrations noted on X-axis). These values are based on measurements of fluorescence intensity in a circular area (50.2µm diameter) near the proximal part of each eye disc (8 discs were used to estimate the mean intensity in each sample).

In order to quantify the levels of mutant polyQ aggregates in eye discs for larvae reared on normal or *KG* or *SG* supplemented food, the immuno-fluorescence intensities of the polyQ protein were measured. Using the maximum intensity projections of confocal z-stack images in the Zen-blue software (Zeiss), the total fluorescence intensities in a circular area (50.2µm diameter) in the proximal region of eye discs immunostained for polyQ aggregates were measured and the means for 8 eye discs for each set were compared. The data, presented in Fig. 3F clearly show that feeding on 0.05% or 0.25% of *KG* or *SG* supplemented food substantially reduced the polyQ aggregates compared to their very high levels seen in eye discs in larvae reared on normal food.

The amyloid aggregates in eye imaginal discs from *ey-GAL4>Aβ42* late 3^rd^ instar larvae reared on normal or *KG* or *SG* supplemented food were immunostained with the anti-Aβ42 antibody. In agreement with the above results on life history parameters, feeding Aβ42 expressing larvae with *KG* or *SG* supplemented food substantially reduced the amyloid accumulation in eye discs (Fig. 4). The higher concentration (0.25%) of *SG* appeared to be slightly less effective than the lower (0.05%).

**Fig. IV.**
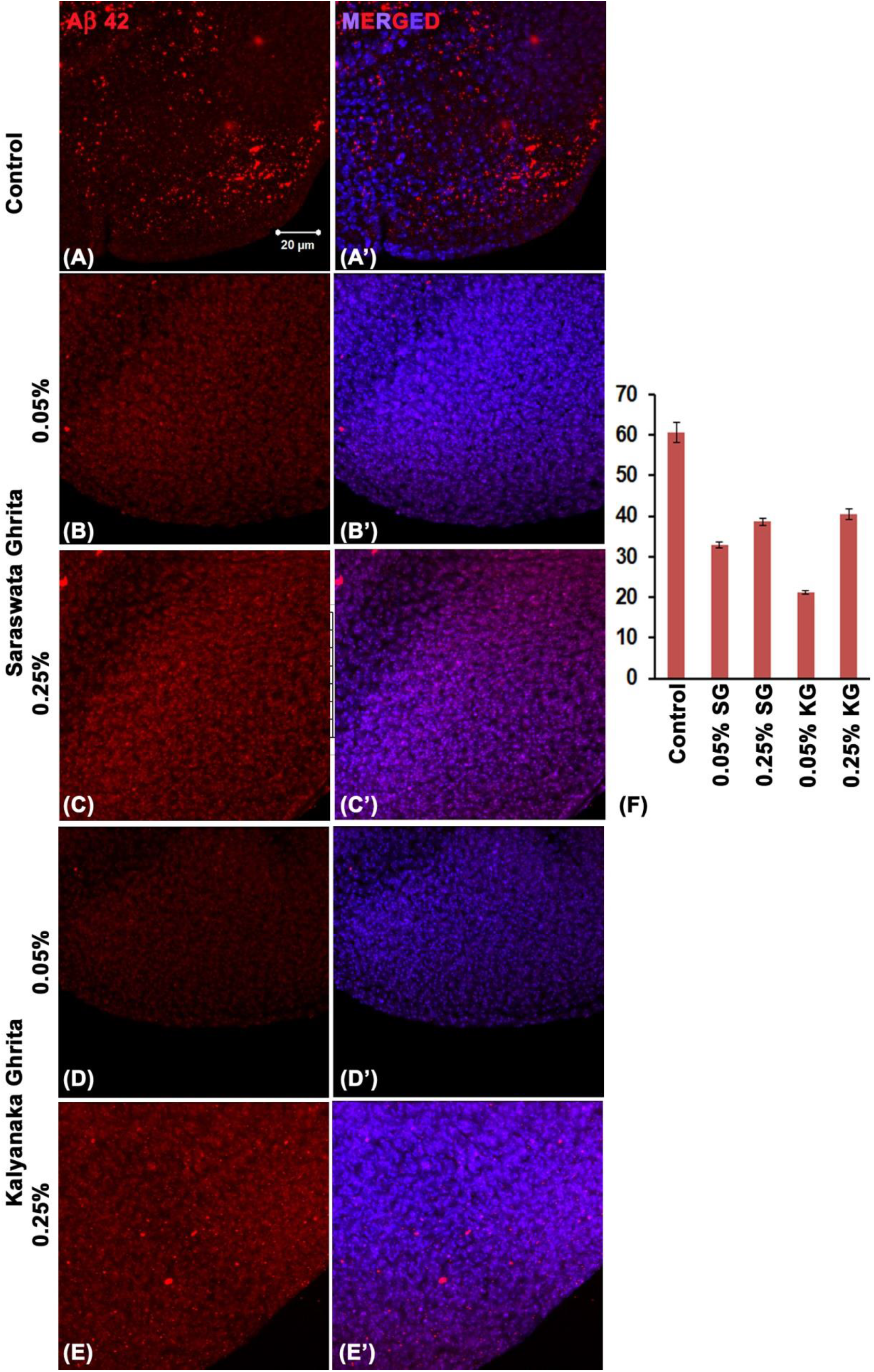
Rearing on *Saraswata Ghrita* or *Kalyanaka Ghrita* supplemented food substantially reduced accumulation of Aβ42 aggregates (red) in *ey-GAL4>UAS-Aβ42* eye discs. (**A-E)** are confocal projection images of *ey-GAL4>UAS-Aβ42* eye discs stained for Aβ42 protein aggregates (red, **A-E**) while **A’-E’** are corresponding merged images including DAPI-stained nuclei (blue). The specific feeding regimes are noted on left side of the rows. (**F**) Histograms of mean (± S.E.) fluorescence intensity of Aβ42 immunostaining (expressed in arbitrary units, Y-axis) in imaginal discs from larvae reared on control or *Saraswata Ghrita* (*SG*) or *Kalyanaka Ghrita* (*KG*) supplemented food (concentrations noted on X-axis). These values are based on measurements of fluorescence intensity in a circular area (50.2µm diameter) near the proximal part of each eye disc (8 discs were used to estimate the mean intensity in each sample).

A quantification of the levels of Aβ aggregates in eye discs from larvae reared on normal or *KG* or *SG* supplemented food, the immuno-fluorescence intensities of the respective proteins were measured as described above for the polyQ expressing discs. The data, presented in Fig. 4F clearly show that feeding on 0.05% or 0.25% of *KG* or *SG* supplemented food substantially reduced the Aβ amyloid plaques compared to their very high levels in eye discs from larvae reared on normal food. The higher concentration (0.25%) of *KG* or *SG* was less effective than their lower concentrations (Fig. 4F).

## 4. Discussion

In this study, we examined effects of dietary administration of two Ayurvedic *ghrita* preparations, viz., *Saraswata* and *Kalyanaka ghritas* on the fruit fly model, using wild type and transgenic strains showing HD and AD symptoms because of targeted expression of respective mutant transgene. Earlier studies in our laboratory (Dwivedi et al 2012, 2013, 2015; Dwivedi and Lakhotia 2016; Tiwari et al 2017, Saba et al, 2017) have established efficacy of dietary administration of Ayurvedic *Rasayanas* in flies as well as mice. When fed at 0.05% (weight/volume) concentration, neither *KG* nor *SG* showed any adverse effects on development of wild type larvae/pupae and lifespan of adult flies, although higher concentrations (0.25% and above) of each of them had dose-dependent adverse effects on life span and/or larval/pupal survival. Therefore, in our studies on the HD and AD flies, we examined effects of 0.05% and 0.25% dietary *KG* or *SG* supplementation.

Classical literature recommends the use of these two *Ayurvedic Ghritas* for treating dementia. Although individual components of the two *ghritas*, especially of the *SG*, have been examined in several studies for their effects on symptoms associated with dementia/AD, there have been very few experimental studies on action or efficacy of the conventional whole formulation. Based on an in-vitro study (Mamidala et al., 2016) on the IMR 32 human neuroblastoma cell line, *SG* was suggested as an effective treatment for ASD (Autism Spectrum Disorder) as it enhanced levels of serotonin, Dopamine and Gamma-aminobutyric acid. In another study (Shelar et al., 2018), *SG* and its lipid extract were found to be effective in reducing Aβ plaques, improve memory and enhance levels of superoxide dismutase, glutathione peroxidase, chloramphenicol acetyl-transferase and nitrous oxide in rat model; interestingly, the lipid extract was found to be more effective than ethanolic extract of *SG*. There does not seem to be any effective experimental study that has examined effects of *KG* or its various component herbs in *in vitro* or *in vivo* models. None of these *ghritas* have so far been examined for action in the context of polyQ toxicity.

Extracts of *Haritaki* (*Terminalia chebula*), a common ingredient in *SG* and *KG*, were shown to protect PC12 cells from Aβ aggregate-mediated cell damages thorough inhibition of ROS and reduction of calcium ion influx (Shen et al 2017). Extract of *Shigru* (*Moringa oleifera*), one of the components of *SG*, was found to defend against anti-oxidant activity and aging in *Cenorhabditis* worm (Chauhan et al 2020; Mahaman et al 2018) and to alleviate tau hyperphosphorylation and Aβ pathology in a homocystein-induced AD rat model. *Moringa oleifera* is also used in Egyptian traditional medicine for improvement of memory and old age-related diseases (Ali et al 2013). Various studies on extracts of different components of *SG*, like *Amalaki, Pippali, Marich* and *Shunthi*, have been reported to inhibit acetylcholinesterase activity, reduce lipid peroxidation in brain, facilitate dissociation of Aβ oligomers, protect against Aβ induced apoptosis, enhance free radical scavenging, chloramphenicol acetyltransferase, superoxide dismutase and glutathione peroxidase activities and thus have beneficial effects in dementia/AD (Farooqui et al., 2018; G et al., 2012; Hritcu et al., 2015; Mahdy et al., 2012; Rashedinia et al., 2021; Saenghong et al., 2012; Sanap et al., 2021; Wiemann et al., 2017; Ali et al., 2013; Chauhan et al., 2020;Tappayuthpijarn et al., 2011; Mathew and Subramanian, 2014; Oboh et al 2012; Ali et al, 2013). In agreement with the earlier studies on *SG* or extracts of its component herbs, our present study on *in vivo* fly model shows that rearing of larvae/flies showing Alzheimer’s neurodegeneration in eyes on food supplemented with 0.05% *SG* reversed the substantially reduced lifespan because of the systemic effects of the toxic protein aggregates. In addition, the present study shows that *KG* is also very effective in suppressing Aβ pathology. In addition, our findings extend the beneficial effects of these two Ayurvedic *ghritas* in polyQ toxicity induced neurodegeneration as well.

It is interesting that while the 0.25% (or even higher, data not presented) dietary concentration of either of the *ghritas* significantly reduced lifespan of wild type flies, the same dosage significantly increased life span of HD or AD flies, although the increase was somewhat less than that seen following feeding on 0.05% *KG* or *SG* supplemented food. This agrees with the *Ayurvedic* principle that dose of a *Rasayana* is specifically related to the individual’s basic health condition (Sharma, 1994). The improved survival of HD or AD transgene expressing organisms following rearing on either of these *ghritas*, correlates with the observed substantial reduction in polyQ or Aβ aggregates in eye discs of respective larvae. Together, these results show, for the first time, especially for the *KG*, that these two *medhyarasayanas* have beneficial effects in *in vivo* AD as well as HD neurodegenerative conditions.

Further studies to understand the molecular bases of actions of these two Ayurvedic *ghrita* formulations will integrate the knowledge based on traditional systems of medicine with the current understanding of cell and molecular biology and thus help in development of better therapies for the increasing burden of age-related dementia and neurodegenerative disorders.

## Acknowledgements

This work was supported by the Distinguished Fellowship grant (no. SB/DF/009/2019) from the Science and Engineering Research Board (Govt of India) to SCL. SS thanks AYUSH Center of Excellence (supported by the Ministry of Ayush, Govt. of India), Center for Complementary and Integrative Health, Interdisciplinary School of Health Sciences, Savitribai Phule Pune University (SPPU), Pune, for fellowship.

## References

Albert, M. S., DeKosky, S. T., Dickson, D., Dubois, B., Feldman, H. H., Fox, N. C., Gamst, A., Holtzman, D. M., Jagust, W. J., Petersen, R. C., Snyder, P. J., Carrillo, M. C., Thies, B., & Phelps, C. H. (2011). The diagnosis of mild cognitive impairment due to Alzheimer’s disease: recommendations from the National Institute on Aging-Alzheimer’s Association workgroups on diagnostic guidelines for Alzheimer’s disease. Alzheimer’s & dementia : the journal of the Alzheimer’s Association, 7(3), 270–279. https://doi.org/10.1016/j.jalz.2011.03.008

Ali, S. K., Hamed, A. R., Soltan, M. M., Hegazy, U. M., Elgorashi, E. E., El-Garf, I. A., & Hussein, A. A. (2013). In-vitro evaluation of selected Egyptian traditional herbal medicines for treatment of Alzheimer disease. BMC complementary and alternative medicine, 13, 121. https://doi.org/10.1186/1472-6882-13-121

Ayurvedic Pharmacopoeia of India, Part-II, Vol.1, p.75&p.85 (1st Ed., 2007), Department of AYUSH, GOI.

Beal, M. F., Brouillet, E., Jenkins, B. G., Ferrante, R. J., Kowall, N. W., Miller, J. M., Storey, E., Srivastava, R., Rosen, B. R., & Hyman, B. T. (1993). Neurochemical and histologic characterization of striatal excitotoxic lesions produced by the mitochondrial toxin 3-nitropropionic acid. The Journal of neuroscience : the official journal of the Society for Neuroscience, 13(10), 4181–4192. https://doi.org/10.1523/JNEUROSCI.13-10-04181.1993

Begum, A. N., Jones, M. R., Lim, G. P., Morihara, T., Kim, P., Heath, D. D., Rock, C. L., Pruitt, M. A., Yang, F., Hudspeth, B., Hu, S., Faull, K. F., Teter, B., Cole, G. M., & Frautschy, S. A. (2008). Curcumin structure-function, bioavailability, and efficacy in models of neuroinflammation and Alzheimer’s disease. The Journal of pharmacology and experimental therapeutics, 326(1), 196–208. https://doi.org/10.1124/jpet.108.137455

Bellen, H. J., Wangler, M. F., & Yamamoto, S. (2019). The fruit fly at the interface of diagnosis and pathogenic mechanisms of rare and common human diseases. Human molecular genetics, 28(R2), R207–R214. https://doi.org/10.1093/hmg/ddz135

Bilen, J., & Bonini, N. M. (2005). Drosophila as a model for human neurodegenerative disease. Annualreviewofgenetics, 39,153–171. https://doi.org/10.1146/annurev.genet.39.110304.095804

Chauhan, A. P., Chaubey, M. G., Patel, S. N., Madamwar, D., & Singh, N. K. (2020). Extension of life span and stress tolerance modulated by DAF-16 in Caenorhabditis elegans under the treatment of Moringa oleifera extract. 3 Biotech, 10(12), 504. https://doi.org/10.1007/s13205-020-02485-x

Chow, C. Y., & Reiter, L. T. (2017). Etiology of Human Genetic Disease on the Fly. Trends in genetics : TIG, 33(6), 391–398. https://doi.org/10.1016/j.tig.2017.03.007

Dwivedi, V., & Lakhotia, S. C. (2016). Ayurvedic Amalaki Rasayana promotes improved stress tolerance and thus has anti-aging effects in Drosophila melanogaster. Journal of biosciences, 41(4), 697–711. https://doi.org/10.1007/s12038-016-9641.

Dwivedi, V., Anandan, E. M., Mony, R. S., Muraleedharan, T. S., Valiathan, M. S., Mutsuddi, M., & Lakhotia, S. C. (2012). In vivo effects of traditional Ayurvedic formulations in Drosophila melanogaster model relate with therapeutic applications. PloS one, 7(5), e37113. https://doi.org/10.1371/journal.pone.0037113

Dwivedi, V, Tripathi B. K. Mutsuddi, M. & Lakhotia, S. C. (2013). Ayurvedic Amalaki Rasayana and Rasa-Sindoor suppress neurodegeneration in fly models of Huntington’s and Alzheimer’s diseases. Curr. Sci. 105, 1711–1723

Dwivedi, V., Tiwary, S., & Lakhotia, S. C. (2015). Suppression of induced but not developmental apoptosis in Drosophila by Ayurvedic Amalaki Rasayana and Rasa-Sindoor. Journal of biosciences, 40(2), 281–297. https://doi.org/10.1007/s12038-015-9521-9

Farooqui, A. A., Farooqui, T., Madan, A., Ong, J. H., & Ong, W. Y. (2018). Ayurvedic Medicine for the Treatment of Dementia: Mechanistic Aspects. Evidence-based complementary and alternative medicine : eCAM, 2018, 2481076. https://doi.org/10.1155/2018/2481076

Finelli, A., Kelkar, A., Song, H. J., Yang, H., & Konsolaki, M. (2004). A model for studying Alzheimer’s Abeta42-induced toxicity in Drosophila melanogaster. Molecular and cellular neurosciences, 26(3), 365–375. https://doi.org/10.1016/j.mcn.2004.03.001

Ganguli, M., Chandra, V., Kamboh, M. I., Johnston, J. M., Dodge, H. H., Thelma, B. K., Juyal, R. C., Pandav, R., Belle, S. H., & DeKosky, S. T. (2000). Apolipoprotein E polymorphism and Alzheimer disease: The Indo-US Cross-National Dementia Study. Archives of neurology, 57(6), 824–830. https://doi.org/10.1001/archneur.57.6.824

Hritcu, L., Noumedem, J. A., Cioanca, O., Hancianu, M., Postu, P., & Mihasan, M. (2015). Anxiolytic and antidepressant profile of the methanolic extract of Piper nigrum fruits in beta-amyloid (1-42) rat model of Alzheimer’s disease. Behavioral and brain functions: BBF, 11, 13. https://doi.org/10.1186/s12993-015-0059-7

Iijima, K., Liu, H. P., Chiang, A. S., Hearn, S. A., Konsolaki, M., & Zhong, Y. (2004). Dissecting the pathological effects of human Abeta40 and Abeta42 in Drosophila: a potential model for Alzheimer’s disease. Proceedings of the National Academy of Sciences of the United States of America, 101(17), 6623–6628. https://doi.org/10.1073/pnas.0400895101

Jameson, J. Larry, 2018 Harrison’s principles of internal medicine, Twentieth edition, Mc-Graw-Hill education, Palatino by Cenveo-Publisher, Pages referred 48, 154, 175&3119.

Jessen, F., Amariglio, R. E., Buckley, R. F., van der Flier, W. M., Han, Y., Molinuevo, J. L., Rabin, L., Rentz, D. M., Rodriguez-Gomez, O., Saykin, A. J., Sikkes, S., Smart, C. M., Wolfsgruber, S., & Wagner, M. (2020). The characterisation of subjective cognitive decline. The Lancet. Neurology, 19(3), 271–278. https://doi.org/10.1016/S1474-4422(19)30368-0

Kumar, S., Harris, R. J., Seal, C. J., & Okello, E. J. (2012). An aqueous extract of Withania somnifera root inhibits amyloid β fibril formation in vitro. Phytotherapy research : PTR, 26(1), 113–117. https://doi.org/10.1002/ptr.3512

Lakhotia, S. C. (2019). Need for integration of Ayurveda with modern biology and medicine. Proc Indian Natn Sci Acad 85, 697–703. https://doi.org/10.16943/ptinsa/2019/49588

Lakhotia, S. C. (2020) Ayurvedic biology-An unbiased approach to revisit and rationalize concepts and practices of Ayurveda. In-Integration Perspectives: Ayurveda, Phytopharmaceuticals and Natural Products (ed. N. Bhat). Continental Prakashan (Pune, India). II/7, pp 87–102

Livingston, G., Huntley, J., Sommerlad, A., Ames, D., Ballard, C., Banerjee, S., Brayne, C., Burns, A., Cohen-Mansfield, J., Cooper, C., Costafreda, S. G., Dias, A., Fox, N., Gitlin, L. N., Howard, R., Kales, H. C., Kivimäki, M., Larson, E. B., Ogunniyi, A., Orgeta, V., … Mukadam, N. (2020). Dementia prevention, intervention, and care: 2020 report of the Lancet Commission. Lancet (London, England), 396(10248), 413–446. https://doi.org/10.1016/S0140-6736(20)30367-6

Mahaman, Y., Huang, F., Wu, M., Wang, Y., Wei, Z., Bao, J., Salissou, M., Ke, D., Wang, Q., Liu, R., Wang, J. Z., Zhang, B., Chen, D., & Wang, X. (2018). Moringa Oleifera Alleviates Homocysteine-Induced Alzheimer’s Disease-Like Pathology and Cognitive Impairments. Journal of Alzheimer’s disease: JAD, 63(3), 1141–1159. https://doi.org/10.3233/JAD-180091

Mahdy, K., Shaker, O., Wafay, H., Nassar, Y., Hassan, H., & Hussein, A. (2012). Effect of some medicinal plant extracts on the oxidative stress status in Alzheimer’s disease induced in rats. European review for medical and pharmacological sciences, 16 Suppl 3, 31–42.

Mallik, M., & Lakhotia, S. C. (2010). Modifiers and mechanisms of multi-system polyglutamine neurodegenerative disorders: lessons from fly models. Journal of genetics, 89(4), 497–526. https://doi.org/10.1007/s12041-010-0072-4

Mamidala, M. P., Rajesh, N., & Rajesh, V. (2016). Mass spectrometric evaluation of neurotransmitter levels in IMR 32 cell line in response to Ayurvedic medicines. Rapid communications in mass spectrometry: RCM, 30(12), 1413–1422. https://doi.org/10.1002/rcm.7571

Mathew, M., & Subramanian, S. (2014). Invitro evaluation of anti-Alzheimer effects of dry ginger (Zingiber officinale Roscoe) extract. Indian journal of experimental biology, 52(6), 606–612.

Matsuda, H., Murakami, T., Kishi, A., & Yoshikawa, M. (2001). Structures of withanosides I, II, III, IV, V, VI, and VII, new withanolide glycosides, from the roots of Indian Withania somnifera DUNAL. and inhibitory activity for tachyphylaxis to clonidine in isolated guinea-pig ileum. Bioorganic & medicinal chemistry, 9(6), 1499–1507. https://doi.org/10.1016/s0968-0896(01)00024-4

McGurk, L., Berson, A., & Bonini, N. M. (2015). Drosophila as an In Vivo Model for Human NeurodegenerativeDisease. Genetics, 201(2),377402.https://doi.org/10.1534/genetics.115.179457

Murthy KRS (ed.); (2009). Astanga Hridaya of Vagbhata. Reprint ed. Chaukhambha Orientalia Publications. 1^st^ Chapter, Uttara Tantra. 14, and 4^th^ Chapter, Uttara Tantra, 44

Nahata, A., Patil, U. K., & Dixit, V. K. (2008). Effect of Convulvulus pluricaulis Choisy. on learning behaviour and memory enhancement activity in rodents. Natural product research, 22(16), 1472–1482. https://doi.org/10.1080/14786410802214199

Nunomura, A., Perry, G., Aliev, G., Hirai, K., Takeda, A., Balraj, E. K., Jones, P. K., Ghanbari, H., Wataya, T., Shimohama, S., Chiba, S., Atwood, C. S., Petersen, R. B., & Smith, M. A. (2001). Oxidative damage is the earliest event in Alzheimer disease. Journal of neuropathology and experimental neurology, 60(8), 759–767. https://doi.org/10.1093/jnen/60.8.759

Oboh, G., Ademiluyi, A. O., & Akinyemi, A. J. (2012). Inhibition of acetylcholinesterase activities and some pro-oxidant induced lipid peroxidation in rat brain by two varieties of ginger (Zingiber officinale). Experimental and toxicologic pathology 64(4), 315–319. https://doi.org/10.1016/j.etp.2010.09.004

Oriel, C., & Lasko, P. (2018). Recent Developments in Using Drosophila as a Model for Human Genetic Disease. International journal of molecular sciences, 19(7), 2041. https://doi.org/10.3390/ijms19072041

Pickett, J., Bird, C., Ballard, C., Banerjee, S., Brayne, C., Cowan, K., Clare, L., Comas-Herrera, A., Corner, L., Daley, S., Knapp, M., Lafortune, L., Livingston, G., Manthorpe, J., Marchant, N., Moriarty, J., Robinson, L., van Lynden, C., Windle, G., Woods, B., … Walton, C. (2018). A roadmap to advance dementia research in prevention, diagnosis, intervention, and care by 2025. International journal of geriatric psychiatry, 33(7), 900–906. https://doi.org/10.1002/gps.4868

Polis, B., & Samson, A. O. (2019). A New Perspective on Alzheimer’s Disease as a Brain Expression of a Complex Metabolic Disorder. In T. Wisniewski (Ed.), Alzheimer’s Disease. Codon Publications.

Rashedinia, M., Mojarad, M., Khodaei, F., Sahragard, A., Khoshnoud, M. J., & Zarshenas, M. M. (2021). The Effect of a Traditional Preparation Containing Piper nigrum L. and Bunium persicum (Boiss.) B.Fedtsch. on Immobility Stress-Induced Memory Loss in Mice. BioMed research international, 2021, 5577594. https://doi.org/10.1155/2021/5577594

Reiter, L. T., Potocki, L., Chien, S., Gribskov, M., & Bier, E. (2001). A systematic analysis of human disease-associated gene sequences in Drosophila melanogaster. Genome research, 11(6), 1114–1125. https://doi.org/10.1101/gr.169101

Rohrer, D., Salmon, D. P., Wixted, J. T., & Paulsen, J. S. (1999). The disparate effects of Alzheimer’s disease and Huntington’s disease on semantic memory. Neuropsychology, 13(3), 381–388. https://doi.org/10.1037//0894-4105.13.3.381

Russo, A., Izzo, A. A., Cardile, V., Borrelli, F., & Vanella, A. (2001). Indian medicinal plants as antiradicals and DNA cleavage protectors. Phytomedicine : international journal of phytotherapy and phytopharmacology, 8(2), 125–132. https://doi.org/10.1078/0944-7113-00021

Saba, K., Rajnala, N., Veeraiah, P., Tiwari, V., Rana, R. K., Lakhotia, S. C., & Patel, A. B. (2017). Energetics of Excitatory and Inhibitory Neurotransmission in Aluminum Chloride Model of Alzheimer’s Disease: Reversal of Behavioral and Metabolic Deficits by Rasa Sindoor. Frontiers in molecular neuroscience, 10, 323. https://doi.org/10.3389/fnmol.2017.00323

Saenghong, N., Wattanathorn, J., Muchimapura, S., Tongun, T., Piyavhatkul, N., Banchonglikitkul, C., & Kajsongkram, T. (2012). Zingiber officinale Improves Cognitive Function of the Middle-Aged Healthy Women. Evidence-based complementary and alternative medicine: eCAM, 2012, 383062. https://doi.org/10.1155/2012/383062

Sanap, A., Joshi, K., Shah, T., Tillu, G., & Bhonde, R. (2021). Pre-conditioning of Mesenchymal Stem Cells with Piper longum L. augments osteogenic differentiation. Journal of ethnopharmacology, 273, 113999. https://doi.org/10.1016/j.jep.2021.113999

Sharma P. V.. (1994). Charaka Samhita of Agnivesha. (Sanskrit with English Translation) 6th edition. Chaukhambha Orientalia, Page no. 132

Shelar, M., Nanaware, S., Arulmozhi, S., Lohidasan, S., & Mahadik, K. (2018). Validation of ethnopharmacology of ayurvedic sarasvata ghrita and comparative evaluation of its neuroprotective effect with modern alcoholic and lipid based extracts in β-amyloid induced memory impairment. https://doi.org/10.1016/j.jep.2018.02.032

Shen, Y. C., Juan, C. W., Lin, C. S., Chen, C. C., & Chang, C. L. (2017). Neuroprotective effect of Terminalia chebula extracts and ellagic acid in pc12 cells. African journal of traditional, complementary, and alternative medicines: AJTCAM, 14(4), 22–30. https://doi.org/10.21010/ajtcam.v14i4.3

Stough, C., Downey, L. A., Lloyd, J., Silber, B., Redman, S., Hutchison, C., Wesnes, K., & Nathan, P. J. (2008). Examining the nootropic effects of a special extract of Bacopa monniera on human cognitive functioning: 90day double-blind placebo-controlled randomized trial. Phytotherapy research: PTR, 22(12), 1629–1634. https://doi.org/10.1002/ptr.2537

Sultana, R., Ravagna, A., Mohmmad-Abdul, H., Calabrese, V., & Butterfield, D. A. (2005). Ferulic acid ethyl ester protects neurons against amyloid beta-peptide(1-42)-induced oxidative stress and neurotoxicity: relationship to antioxidant activity. Journal of neurochemistry, 92(4), 749–758. https://doi.org/10.1111/j.1471-4159.2004.02899.x

Tappayuthpijarn, P., Itharat, A., & Makchuchit, S. (2011). Acetylcholinesterase inhibitory activity of Thai traditional nootropic remedy and its herbal ingredients. Journal of the Medical Association of Thailand = Chotmaihet thangphaet, 94 Suppl 7, S183–S189.

Tiwari, V., Saba, K., Veeraiah, P., Jose, J., Lakhotia, S. C., & Patel, A. B. (2017). Amalaki Rasayana improved memory and neuronal metabolic activity in AbPP-PS1 mouse model of Alzheimer’s disease. Journal of biosciences, 42(3), 363–371. https://doi.org/10.1007/s12038-017-9692-7

Uabundit, N., Wattanathorn, J., Mucimapura, S., & Ingkaninan, K. (2010). Cognitive enhancement and neuroprotective effects of Bacopa monnieri in Alzheimer’s disease model. Journal of ethnopharmacology, 127(1), 26–31. https://doi.org/10.1016/j.jep.2009.09.056

Wiemann, J., Karasch, J., Loesche, A., Heller, L., Brandt, W., & Csuk, R. (2017). Piperlongumine B and analogs are promising and selective inhibitors for acetylcholinesterase. European journal of medicinal chemistry, 139, 222–231. https://doi.org/10.1016/j.ejmech.2017.07.081.

Zoghbi, H. Y., & Orr, H. T. (2000). Glutamine repeats and neurodegeneration. Annual review of neuroscience, 23, 217–247. https://doi.org/10.1146/annurev.neuro.23.1.217

